# Symbiont biosynthetic output tracks host developmental stage in a marine sponge holobiont

**DOI:** 10.64898/2026.09.29.755549

**Authors:** Eve A. Maunders, Zhee Sheen Wong, Rebecca Fieth, Xueyan Xiang, Bernard M. Degnan, Sandie M. Degnan

**Author notes:** Address correspondence to Sandie M. Degnan. These authors contributed equally to this work (Eve A. Maunders, Zhee Sheen Wong).

## Abstract

Microbes have likely influenced animal development since its origin, yet how host and microbe partner across ontogeny remains poorly understood. Here we characterise a developmentally dynamic symbiosis in the marine sponge *Amphimedon queenslandica* by simultaneously profiling host and symbiont gene expression through embryonic, larval and metamorphic development. The transcriptional states of all three core, vertically inherited symbionts were strikingly stage-specific. Metabolic gene expression of the dominant symbiont, *AqS1*, peaked in brooded embryos, when host glycolytic and oxidative pentose phosphate pathway expression was lowest, and this peak was enriched for enzyme-encoding genes. In non-feeding larvae, *AqS1* biosynthetic gene expression contracted across central carbon, vitamin and amino acid metabolism, while its import capacity increased. This contraction was selective. *AqS1* genes for the final steps of arginine synthesis, which the host lacks, were minimally expressed in pre-competent larvae and upregulated at settlement together with host nitric oxide synthase, consistent with the known requirement for symbiont-derived arginine in nitric oxide-induced settlement. Histidine biosynthesis genes were instead maintained in the larva, coinciding with upregulation of host histidine-to-glutamate conversion. Host and symbiont nutrient transporters peaked in the same developmental stages more often than expected by chance. In the filter-feeding juvenile, host metabolism and efflux transport reached their maximum while the symbiont transcriptome changed little. These findings show that symbiont biosynthetic output tracks host developmental stage, and suggest that an inherited microbiome may contribute to successive stages of an animal life cycle through changes in its transcriptional activity rather than its composition.

**Importance statement:** Animals develop in partnership with microbes, but how symbionts contribute across distinct stages of a host life cycle is poorly understood. We followed a vertically inherited microbiome through the development of a marine sponge, a member of one of the earliest-branching animal lineages, from brooded embryo to feeding juvenile. The dominant symbiont did not simply grow or decline with its host, but rather it reprogrammed its metabolism at each developmental stage, including the timing of its arginine synthesis, which the host requires to produce the nitric oxide signal for settlement. Our study shows that transcriptional reprogramming, rather than changes in community composition, is one way inherited microbes may shape animal development, and suggests that such developmental integration may be an ancient feature of animal–microbe symbiosis.

## Introduction

Microbial symbionts are ubiquitous in all major animal lineages and often persist throughout the host life cycle (Carrier & Bosch, 2022). Indeed, the first multicellular animals likely comprised not only their own zygotically-derived cells but also associated microbes (Bosch & Hadfield, 2020; Gilbert & Hadfield, 2022; Margulis, 1991; McFall-Ngai et al., 2013). Over deep evolutionary time, this intimacy has produced a spectrum of dependencies, among the most profound of which are developmental symbioses in which normal animal ontogeny requires specific microbial signals (Gilbert, 2016). Such interdependences span diverse taxa. For example, the Hawaiian bobtail squid requires luminescent *Vibrio fischeri* symbionts to drive remodeling of its light organ (Nyholm & McFall-Ngai, 2021); bacterial colonisation of the mammalian gut during infancy shapes immune development (Round et al., 2011; Sawa et al., 2011; Tanaka & Nakayama, 2017) and can influence life history tradeoffs (Petrullo et al., 2026); symbionts influence insect development (Kang et al., 2021; Mirth et al., 2019); and in marine sponges, symbionts are digested to provision yolk during embryogenesis in *Halichondria panicea* (Turon et al., 2026) and to supply arginine required for larval settlement in *Amphimedon queenslandica* (Song et al., 2021).

In adult animal holobionts, nutrient exchange is a well-known hallmark of symbiosis. Hosts provide stable, nutrient-rich habitats, while symbionts expand host metabolic diversity by fixing carbon and nitrogen, and synthesising essential vitamins and amino acids that the host cannot produce (Burriesci et al., 2012; Feng et al., 2019; Hentschel et al., 2012; Muscatine et al., 1984; Rix et al., 2020; Song et al., 2021; Zahran, 1999; Zhang et al., 2024). Much less is known about how these metabolic partnerships change as the host itself changes. Animal physiology, nutrition and immune status shift dramatically through development, so that metabolic contracts between host and symbionts must also shift. This is particularly striking in animals with biphasic life cycles that alternate between pelagic and benthic phases, a pattern likely ancestral for metazoans (Boyle & Young, 2025; Degnan & Degnan, 2023). The drastic ecological and physiological transitions between phases provide a natural framework for investigating how symbiont functions adjust to developmental change.

The marine demosponge *Amphimedon queenslandica* hosts a low complexity-low abundance symbiont community dominated by three extracellular, heterotrophic bacterial symbionts (hereafter *AqS1*, *AqS2* and *AqS3*) that are maternally-inherited; *AqS1*, a sulfur-oxidizing gammaproteobacterium of the order Chromatiales, dominates the microbiome throughout the life cycle (Figure 1) (Fieth et al., 2016; Gauthier et al., 2016). Because they are transmitted early in embryogenesis (Fieth et al., 2016), these symbionts are present at every stage of the life cycle and thus have potential to influence host development. As a consequence of the pelagobenthic life cycle, each stage occupies a distinct habitat: embryos are brooded within the benthic adult, larvae swim freely in the plankton, and the sessile postlarvae, juveniles and adults are attached to the benthos, with the juvenile marking the beginning of active filter-feeding on environmental bacteria. Prior work from our group has shown that these symbionts are essential for host larval settlement and metamorphosis: they synthesise arginine, which the larva uses as the obligate substrate for nitric oxide (NO) synthesis, and NO is required for the induction of settlement (Ueda et al., 2016; Song et al., 2021). This raises the question of how the broader metabolic relationship between host and symbiont changes across development, and how the timing of symbiont-derived arginine fits within it.

**Figure 1.**
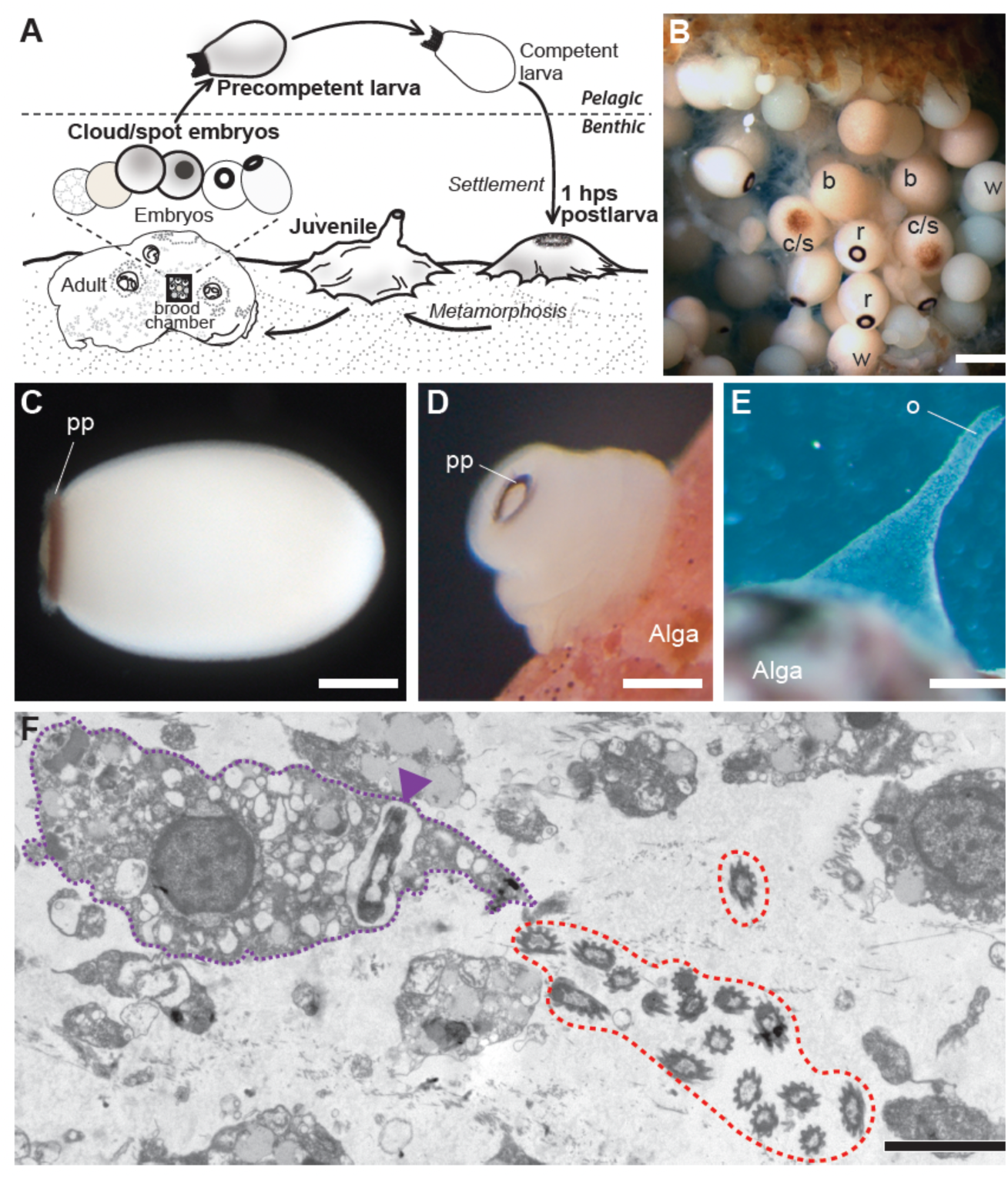
Life cycle of the *Amphimedon queenslandica* holobiont. (A) Schematic depicting the biphasic life cycle of the sponge host, with pelagic and benthic stages separated by the horizontal dashed line. Developmental stages analysed in this study are in bold: cloud/spot stage embryos; precompetent larva; 1 hour post-settlement (hps) postlarva; and 4 day old juvenile (adapted from Yuan et al., 2022). (B) Brood chamber of an adult sponge with white (w), brown (b), cloud/spot (c/s) and ring (r) embryonic stages labelled; bacterial symbionts are transferred into white and brown stage embryos via adult nurse cells (Fieth et al., 2016). (C) Larval stage showing the posterior pigment ring (pp). (D) Early metamorphosing postlarval stage settled on coralline alga (0.5-1 hps) and with the pp still visible. (E) Juvenile stage with an exhalant osculum (o). (F) Transmission electron micrograph, generated as in Fieth et al. (2016), showing several *AqS1* inhabiting extracellular space in the juvenile sponge mesohyl (circled by red dashed line), and a single *AqS1* (purple arrow) phagocytosed by a sponge cell (outlined by purple dotted line). Scale bars: (B) 500 µm; (C-E) 100 µm; (F) 1 µm.

To address this question, we compare host and symbiont transcriptomes across four developmental stages – embryos, swimming larvae, newly settled postlarvae and sessile juveniles (Figure 1A-E) – using a dual RNA-Seq approach that profiles both partners simultaneously. Parallel host and symbiont transcriptomics has recently been applied across the adult reproductive cycle of *H. panicea*, revealing phagocytosis of symbionts during early embryogenesis (Turon et al., 2026). Here we instead follow discrete developmental stages of the offspring, in a holobiont with partner-resolved genomes (Gauthier et al., 2016; Xiang et al., 2026), to trace symbiont contributions through ontogeny at the level of individual pathways and enzymatic steps.

We find that symbiont biosynthetic output tracks with host developmental stage: it peaks in embryogenesis, contracts during the non-feeding larval phase, and increases again at settlement. Within the larval contraction, however, specific amino acid pathways are regulated independently. Symbiont arginine biosynthesis is reduced in the larva and restored at settlement, consistent at the transcriptional level with the experimental demonstration that symbiont-derived arginine is required for settlement (Song et al., 2021), while histidine biosynthesis is maintained in the larva, coinciding with host conversion of histidine to glutamate. Sponges are one of the earliest-branching animal lineages (Schultz et al., 2023; Simion et al., 2017; Srivastava et al., 2010) with an ancestral-type biphasic life cycle, and the presence of a developmental symbiosis in this system therefore bears on whether integration between an animal and its inherited microbiome across the life cycle was a feature of the last common ancestor of extant animals.

## Results

### Host and symbiont transcriptomes are stage-specific across sponge development

Having previously identified the metabolic potential of core bacterial symbionts within the *A. queenslandica* hologenome and adult holotranscriptome (Gauthier et al., 2016; Xiang et al., 2026), here we focussed on how expression of metabolism-related genes across holobiont partners changes through development. We used a dual RNA-Seq approach to simultaneously capture biologically-replicated transcriptomes from both host and bacteria from four developmental stages: (i) cloud-spot embryos in maternal brood chambers, (ii) swimming pre-competent larvae, (iii) postlarvae one hour after settlement (1 hps), and (iv) four day old filter-feeding juveniles (Figure 1). Raw reads are available at NCBI under Bioproject PRJNA982967.

Each stage yielded an average of >135 million quality-filtered reads, and most of the aligned reads (98.9%) mapped to *A. queenslandica*, with 1.10, 0.012 and 0.032% mapping to *AqS1, AqS2* and *AqS3*, respectively (Supplementary Table 1). Mean protein-coding sequence (CDS) coverage across the developmental holotranscriptomes was 81.9% for *A. queenslandica*, 89.9% for *AqS1*, 35.0% for *AqS2*, and 40.5% for *AqS3*. Median read depth per gene was high for *A. queenslandica* (57) and *AqS1* (157), but low for *AqS2* and *AqS3* (3). This ∼50-fold difference in read depth relative to *AqS1* means that our consideration of differential expression for *AqS2* and *AqS3* is confined to their most highly expressed genes (upper quartiles Q3–Q4). Count tables are available as Supplementary Table 2.

The transcriptomes of each holobiont partner were strikingly stage-specific across developmental stages (Figure 2). Among the bacterial symbionts, the transcriptomes of benthic stages (1 hps postlarvae and juveniles) were most similar to each other, while embryos and the pelagic larval stage were more divergent. In contrast, the four developmental transcriptomes of the sponge host were roughly equidistant from one another. Because the developmental transcriptome of the host *A. queenslandica* has been characterised previously across the life cycle (Conaco et al., 2012; Fernandez-Valverde et al., 2015; Levin et al., 2016) and in detail through larval competence, settlement and early metamorphosis (Say & Degnan, 2020; Yuan et al., 2026), we focus here on the symbiont transcriptomes and on the host genes that interact directly with symbiont-derived metabolites.

**Figure 2.**
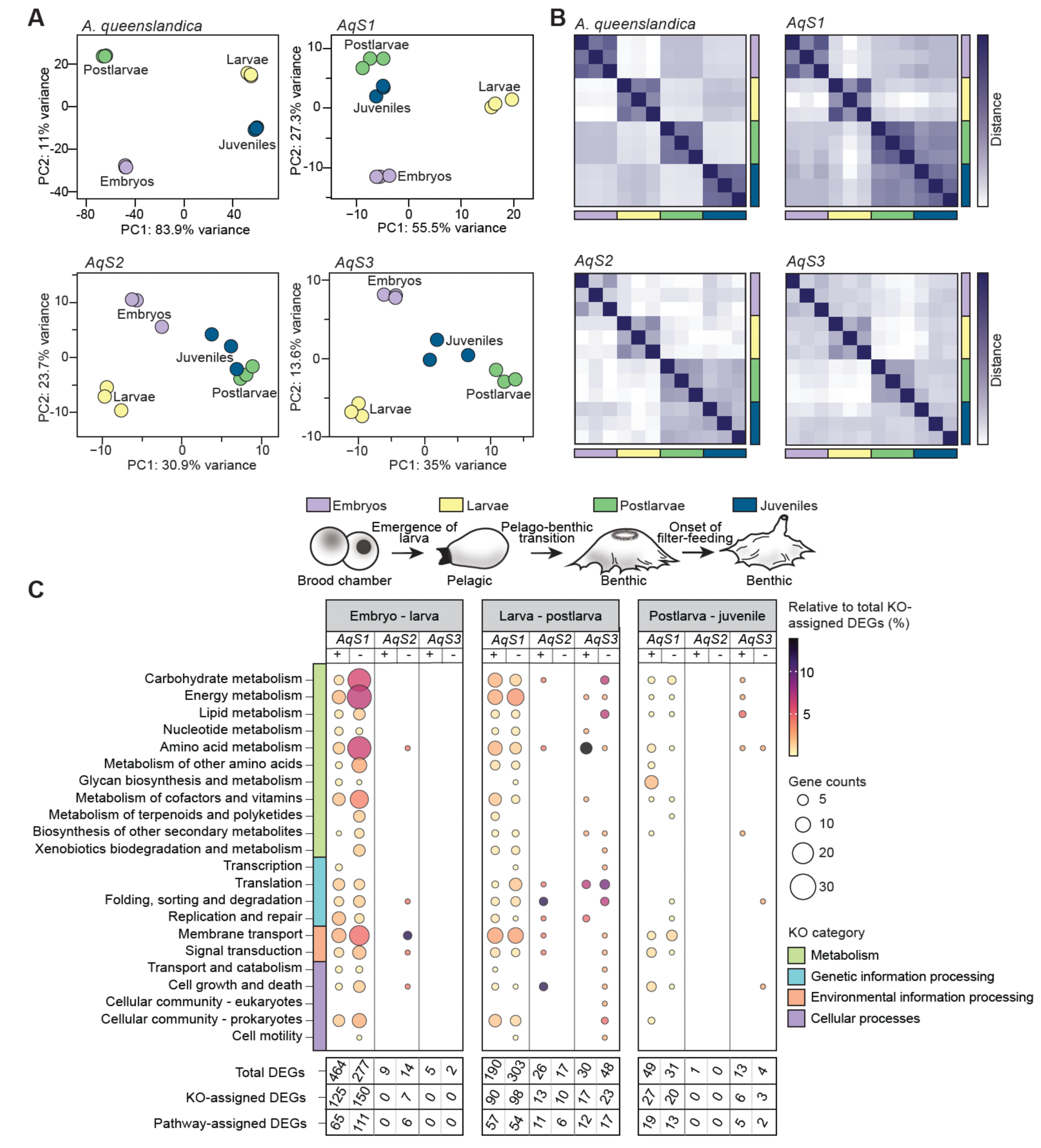
Stage-specific gene expression in the *A. queenslandica* holobiont. (A) Principal component analysis of holotranscriptome profiles across four developmental stages. Embryonic (purple), larval (yellow), 1 hps postlarval (green) and 4 day old juvenile (blue) stages shown. (B) Heatmaps of biologically replicated developmental transcriptomes showing pairwise distances based on gene expression, calculated using Euclidean metrics. Dark purple indicates high similarity (minimum distance = 0), while white indicates low similarity (maximum distance, corresponding to the largest calculated pairwise distance). Replicates from low read-coverage samples (*AqS2* and *AqS3*) show greater pairwise distances, possibly reflecting increased noise and gene-level dropout associated with low sequencing depth. (C) KEGG pathways represented by differentially expressed genes in bacterial symbionts across the four stages. Differentially expressed genes (DEGs) from symbionts *AqS1*, *AqS2* and *AqS3* were mapped to KEGG pathways. Dot size indicates the number of significantly upregulated (+) and downregulated (-) genes assigned to each pathway. Dot colour reflects the relative representation of each pathway, calculated as the proportion of pathway-assigned genes relative to the total number of KEGG orthology (KO)-assigned DEGs in the dataset for each symbiont (*AqS1*, 396; *AqS2*, 25; *AqS3*, 41). Read count data and differential expression statistics used to generate these figures are available in Supplementary Tables 2 and 3, respectively.

Pairwise differential expression analyses were performed on sequential developmental stages (Supplementary Table 3). In total, 1,127 symbiont genes were significantly differentially expressed (using criteria |log2FC| ≥1 and padj ≤0.05) in at least one pairwise comparison, of which 87.8% were encoded by *AqS1*, 4.7% by *AqS2* and 7.5% by *AqS3*. The largest transcriptional shifts for the symbionts occurred from cloud/spot embryonic to competent larval stages, with a total of 478 symbiont genes upregulated and 293 downregulated, and from competent larval to 1 hps postlarval stages (245 symbiont genes upregulated and 368 downregulated) (Supplementary Figure 1). In contrast, the transition between benthic stages (from 1 hps postlarvae to juveniles, which includes the onset of filter feeding) showed the fewest changes in symbiont transcriptomes with 63 and 35 genes being up- and downregulated, respectively.

Of the 1,127 symbiont differentially expressed genes (DEGs), 462 were KEGG-annotated; 303 of these mapped to specific KEGG Pathways (Figure 2C; Supplementary Table 3). These represented 25.8%, 34.0%, and 35.7% of DEGs from *AqS1*, *AqS2*, and *AqS3*, respectively. Metabolism-related pathways, particularly carbohydrate, energy, and amino acid metabolism were the most highly represented pathways among the DEGs, with 91.0% of these metabolic DEGs encoded by *AqS1*. The ‘environmental information processing’ category contained the second largest set of DEGs, with ‘membrane transport’ genes particularly well represented. *AqS1* again accounted for the majority (86.5%) of genes in this category, having numerous ABC transporters involved in translocation of metabolic substrates, such as amino acids, sugars and phosphate that were dynamically expressed across development. The direction of these changes differed by transition (Figure 2C). At the embryo-to-larva transition, *AqS1* upregulated more genes than it downregulated overall (464 versus 277), but among pathway-assigned DEGs the balance reversed (65 upregulated versus 111 downregulated), and the downregulated set was dominated by carbohydrate, energy, amino acid and cofactor/vitamin metabolism. The embryo-to-larva transition thus combined a broad upregulation of largely unannotated genes with a specific contraction of metabolic pathway expression.

### Symbiont metabolic gene expression peaks during embryogenesis

The most striking shifts in metabolic gene expression that were observed across *A. queenslandica* development involved central carbon metabolism, including glycolysis, the tricarboxylic acid (TCA) cycle, and the pentose phosphate pathway (PPP). The host sponge *A. queenslandica* encodes complete glycolysis, TCA cycle and PPP pathways, all of which were highly expressed (quartile 4; Q4) throughout development (Figure 3A), although glycolytic and oxidative PPP genes reached their highest expression in juveniles (Figure 3B).

**Figure 3.**
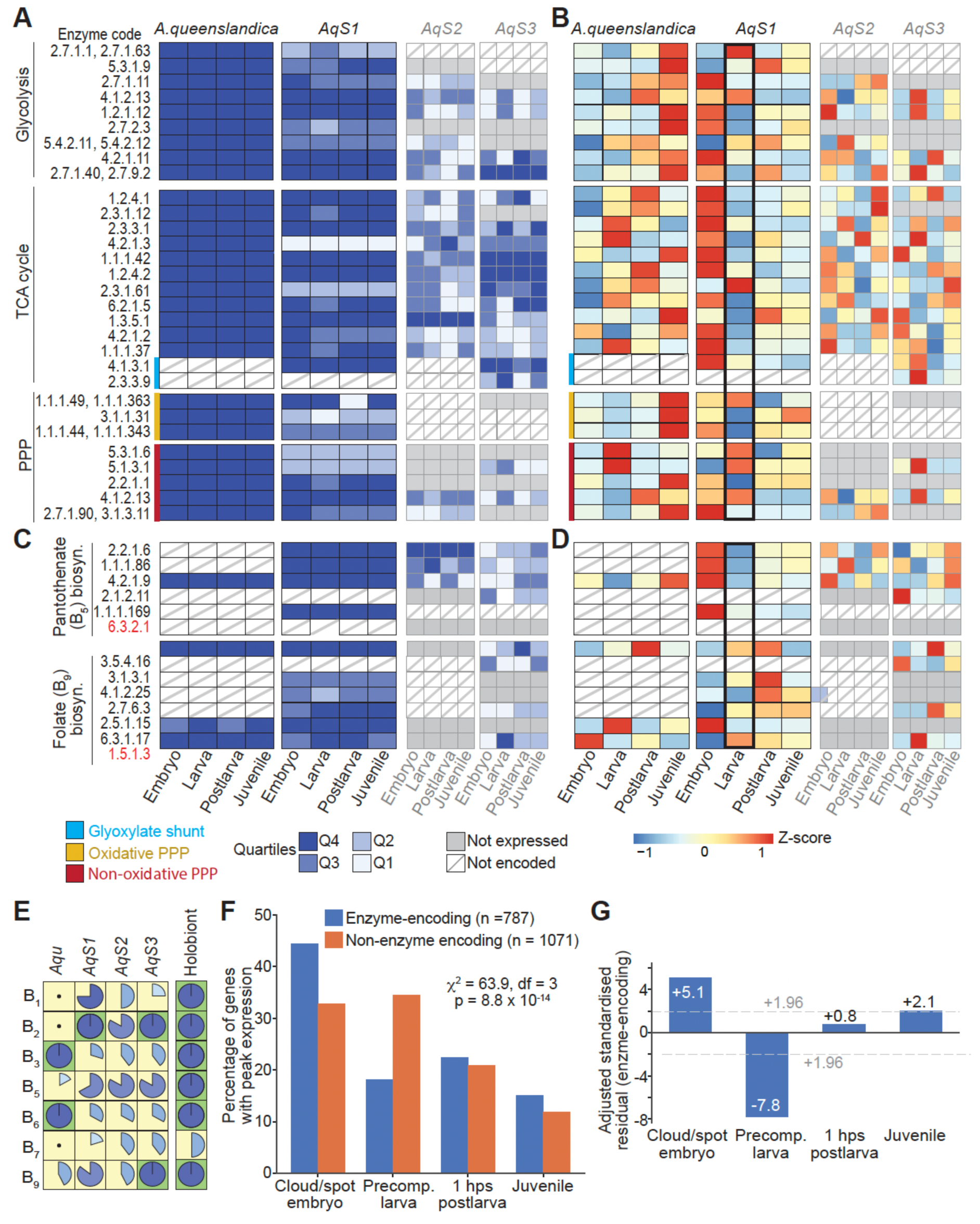
*AqS1* metabolism is high in embryos then contracts in larvae. (A,B) Expression levels of genes involved in energy production and carbohydrate metabolism in the host and three symbionts (*AqS1*, *AqS2* and *AqS3*) across four developmental stages, shown as (A) quartiles, with Q4 being the top 25% of expressed genes within each transcriptome, and (B) Z-scores normalised individually for each holobiont partner per gene. *AqS1* larval expression (black border) shows a pattern of metabolic constraint across carbon metabolism pathways. (C,D) Expression levels of B_5_ and B_9_ vitamin biosynthesis pathways, which are partially encoded by the host, shown as (C) quartiles and (D) Z-scores. Enzyme codes coloured in red represent the final enzyme in the biosynthesis pathway. (E) Vitamin biosynthesis capacity across holobiont partners shown by percent pathway completeness. Mean normalised count, quartile and EC data used to generate A-E are available in Supplementary Table 4. (F,G) The embryonic peak of *AqS1* transcription is enriched in enzyme-encoding genes. (F) Percentage of enzyme-encoding (EC-annotated; blue, n = 787) and non-enzymatic (orange, n = 1,071) *AqS1* genes with peak expression at each developmental stage. Only genes expressed in the upper two quartiles (Q3 or Q4) in at least one stage are included. The peak-stage distributions differ between the two classes (χ² = 63.9, df = 3, p = 8.8 × 10⁻¹⁴). (G) Adjusted standardised residuals for enzyme-encoding genes at each stage; residuals for non-enzymatic genes are equal in magnitude and opposite in sign. Dashed lines at ±1.96 mark departures significant at α = 0.05. Data used to generate F,G are available in Supplementary Table 5.

*AqS1* likewise encodes complete pathways for glycolysis, TCA cycle and PPP, with most enzymes expressed at moderate to high levels (Q3–Q4) throughout development (Figure 3A). However, in contrast to the host, *AqS1* glycolytic and TCA cycle gene expression was highest in embryos and declined markedly in larvae before partially recovering in post-settlement stages (Figure 3B). Of the symbionts, only *AqS1* encodes and expressed the complete PPP oxidative branch, though at lower levels than the host (*AqS1*: Q2–Q4; host: Q4; Figure 3A). During embryogenesis, when host oxidative PPP activity was relatively low, *AqS1* expression of the oxidative branch was at its peak (Figure 3B).

The picture of symbiont metabolic support also encompassed vitamin biosynthesis. *A. queenslandica* is missing the biosynthetic pathways for B₁, B₂ and B₇ entirely and encodes only partial pathways for vitamins B₅ and B₉, so that symbionts are required for *de novo* synthesis of these B vitamins (Xiang et al., 2026; Figure 3E). Vitamin B₅ (pantothenate) is the obligate precursor for coenzyme A (CoA), a central cofactor in energy metabolism, fatty acid synthesis and the TCA cycle (Czumaj et al., 2020). *A. queenslandica* encodes only one of the six enzymes required for pantothenate synthesis, whereas *AqS1* encodes four, and all were expressed at high levels (Q4) throughout development (Figure 3C). Strikingly, the *AqS1* pantothenate biosynthesis expression profile closely mirrored that of its TCA cycle genes, with peak expression in embryos, a marked decline in larvae and partial recovery in postlarva (Figure 3D). Interestingly, the remaining two genes required for completion of the pantothenate synthesis pathway, including the final step (EC 6.3.2.1), are encoded by *AqS2* and *AqS3*. In the present study, transcripts for that final step were not detected in either symbiont, potentially because of low sequencing depth, so the terminal conversion remains unaccounted for at the transcriptional level.

To determine whether the embryonic peak in *AqS1* metabolic gene expression reflects a general elevation of transcription at this stage or a specific concentration of enzymatic function, we compared the developmental timing of enzyme-encoding (EC-annotated) and non-enzymatic *AqS1* genes (Supplementary Table 5). The comparison was restricted to genes expressed in Q3 or Q4 in at least one stage (787 enzyme-encoding, 1,071 non-enzymatic). The two gene classes differed in their distribution of peak stages (χ² = 63.9, df = 3, p = 8.8 × 10⁻¹⁴; Figure 3F,G). Enzyme-encoding genes were over-represented among embryo-peaking genes (44.3% versus 32.8% of non-enzymatic genes; odds ratio 1.63, 95% CI 1.35–1.98; adjusted standardised residual +5.1) and under-represented among larva-peaking genes (18.2% versus 34.5%; residual −7.8). There was no difference for postlarva (22.4% versus 20.9%) or juvenile (15.1% versus 11.9%) (Figure 3F,G). Non-enzymatic genes were, by contrast, as likely to peak in larvae as in embryos, and two-thirds (66%) of those peaking in larvae were annotated as hypothetical or uncharacterised proteins, compared with half (50%) of those peaking in embryos. Thus the high concentration and expression of *AqS1* metabolic genes in the embryo is not simply part of a general elevation in transcriptional activity at this life stage, but appears instead to indicate a specific concentration of enzymatic gene expression in the embryo.

### Symbiont biosynthesis selectively contracts in the larval stage

The embryo-to-larva transition produced the largest transcriptional shift in the symbionts (Supplementary Figure 1, Supplementary Table 3), and its dominant feature was a contraction of *AqS1* biosynthetic gene expression that extended across central carbon metabolism (Figure 3A,B), B-vitamin synthesis (Figure 3C,D) and amino acid synthesis (Figure 4, Supplementary Figure 2, Supplementary Table 4). This contraction was not uniform, however, and the pathways that departed from it are described here alongside the contraction.

**Figure 4.**
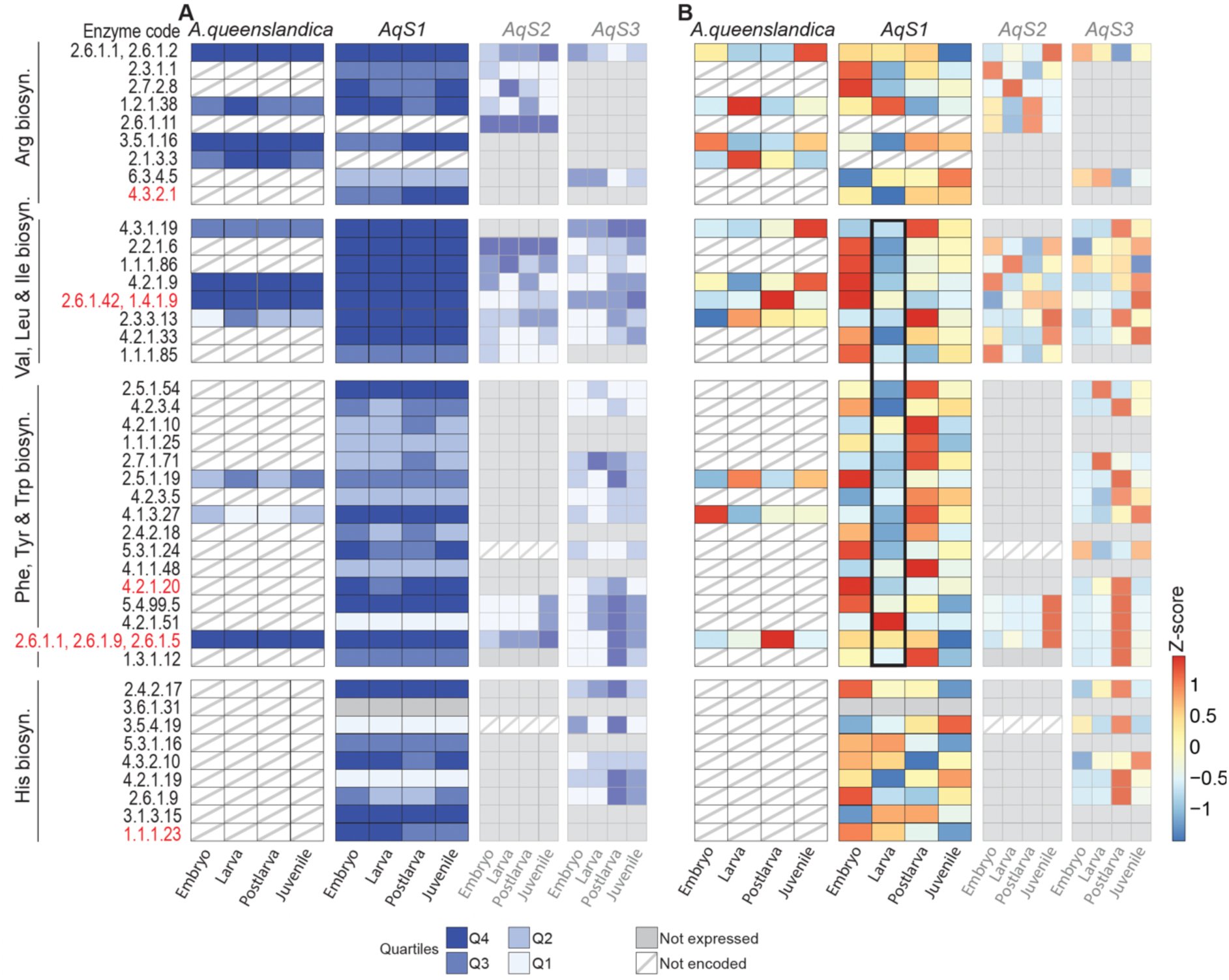
Expression of amino acid biosynthesis genes in *A. queenslandica* and its bacterial symbionts. Transcript levels of genes involved in amino acid biosynthesis in the host and symbionts (*AqS1*, *AqS2* and *AqS3*) across four developmental stages are shown as (A) quartiles, with Q4 representing the top 25% of expressed genes within each transcriptome, and (B) Z-scores normalised individually for each holobiont partner, per gene. Enzyme codes coloured red denote the final enzyme in the biosynthesis pathway. *AqS1* larval expression (black border) shows a pattern of metabolic constraint across amino acid biosynthesis pathways, except arginine and histidine. Mean normalised count, quartile and EC data used to generate this figure are available in Supplementary Table 4

The *A. queenslandica* genome lacks complete *de novo* biosynthetic pathways for eleven amino acids, while the combined genetic repertoire of its three symbionts complements these deficiencies (Xiang et al., 2026). A notable feature of this partitioning is that the host encodes the final enzymatic step in the synthesis of isoleucine, leucine, valine, methionine, phenylalanine, tyrosine and lysine, but lacks the upstream intermediary enzymes (Xiang et al., 2026; Figure 4, Supplementary Figure 2).

Throughout development, *AqS1* carried the majority of the amino acid biosynthetic load, expressing genes for most of the host-deficient pathways at moderate to high levels (Figure 4A). In pre-competent larvae, *AqS1* expression of genes for lysine, valine, leucine, isoleucine, phenylalanine, tyrosine, tryptophan and methionine synthesis was markedly reduced (Figure 4B, Supplementary Figure 2), matching the contraction of central carbon and vitamin metabolism reported above, and coinciding with the stage at which the non-feeding holobiont faces its greatest nutritional challenge. These pathways were upregulated again in 1 hps postlarvae, in concert with the upregulation of the host’s terminal enzymes (Figure 4, Supplementary Table 3).

Two amino acid pathways did not follow this pattern, histidine and arginine (Figure 4; Supplementary Table 4). The two, sequential terminal steps of *AqS1* histidine biosynthesis (EC 3.1.3.15, 1.1.1.23) were maintained at high levels through the larval phase despite the decline in several upstream steps, and fell only after settlement. Conversely, the citrulline-to-arginine module of *AqS1* arginine biosynthesis, argininosuccinate synthase (EC 6.3.4.5) and argininosuccinate lyase (EC 4.3.2.1), did not peak in embryos as in the other pathways and was minimally expressed in larvae. Instead, it was upregulated around the time of settlement. Both exceptions concern precursors for host signalling molecules and are examined in the following section.

Sulfur energy metabolism was likewise selectively regulated (Supplementary Figure 2; Supplementary Table 4). *AqS1* sulfur oxidation (*soxAXYZ*) genes were highly expressed throughout development, whereas the ATP-generating adenylylsulfate reductase and sulfate adenylyltransferase genes (*aprAB*, *sat*) of the reverse dissimilatory sulfate reduction pathway (Loy et al., 2009) followed the general biosynthetic pattern, declining from embryo to larva and recovering at settlement, while the dissimilatory sulfite reductase genes (*dsrABEFH*) of the same pathway peaked in the larva. The larval peak of the Dsr module was thus decoupled from the AprAB/Sat module of the same pathway.

Finally, the larval contraction of biosynthesis was accompanied by upregulation of *AqS1* import capacity. Among *AqS1* differentially expressed transporters, a distinct cluster peaked in the larva, comprising sugar (*rbs*), branched-chain amino acid (*liv*) and oligopeptide (*opp*) ABC transporter components (Supplementary Figure 3; Supplementary Table 7). *AqS1* also encodes a ten-paralog family of C4-dicarboxylate TRAP transporter permease genes (*dctM*), of which the majority of the six differentially expressed genes were upregulated from embryo to larva (Supplementary Table 3), consistent with increased uptake of TCA-cycle intermediates such as malate, fumarate and succinate as alternative carbon sources during the period of biosynthetic restriction. The larval *AqS1* transcriptome therefore reflects a shift from synthesis toward acquisition of host-derived substrates, with biosynthesis retained only in specific pathways.

### Arginine and histidine biosynthesis depart from the larval contraction

The two *AqS1* amino acid pathways that departed from the larval contraction may be relevant to signalling as well as nutrition. The arginine pathway is linked by experimental evidence to settlement (Song et al., 2021), and the histidine pathway feeds a host glutamate pathway implicated in larval phototactic behaviour (Wong et al., 2022). In each case the symbiont’s biosynthetic gene expression rises and falls in parallel with the host genes that use its products.

#### Arginine and nitric oxide

Arginine biosynthesis requires the combined contributions of *AqS1* and either *AqS2* or *AqS3*, and its final step of argininosuccinate lyase (ASL; EC 4.3.2.1) is absent from the host genome. The host does, however, constitutively express the citrulline-forming step, ornithine carbamoyltransferase (EC 2.1.3.3), at high levels throughout development (Figure 4). Within *AqS1,* the pathway was not uniformly expressed. The upstream N-acetylglutamate steps (EC 2.3.1.1, 2.7.2.8) followed the embryonic peak and larval trough characteristic of *AqS1* metabolism generally, whereas the ornithine-releasing step (EC 3.5.1.16) and the citrulline-to-arginine module (ASS; EC 6.3.4.5, ASL) lacked the embryonic peak and were induced at settlement, ASL rising from its lowest levels in embryos and pre-competent larvae to high expression in 1 hps postlarvae and juveniles (Figure 4; Supplementary Table 4). The host matched this expression profile from the other side: host nitric oxide synthase (NOS; EC 1.14.13.39), which converts arginine to NO, increased progressively from embryos to larvae and from larvae to postlarvae before declining in juveniles. By contrast, *AqS1* nitric oxide reductase (NOR; EC 1.7.2.5), which degrades NO to nitrous oxide, was most highly expressed in pre-competent larvae, the stage that lacks the capacity to settle (Supplementary Figure 2; Supplementary Table 4). A further host–symbiont pairing supported this settlement axis, with both the bacterial-type high-affinity zinc uptake system of *AqS1* (*znuA*, *znuB*, *znuC*) peaking in postlarvae (Supplementary Table 7), and the host’s zinc efflux transporter, solute carrier family 30 member 2, being strongly upregulated at the same stage before declining in juveniles (Supplementary Table 6). A bacterial zinc importer and a host zinc exporter peaking together at settlement is consistent with directional host-to-symbiont zinc transfer at this stage.

#### Histidine and glutamate

*A. queenslandica* does not encode histidine biosynthetic enzymes but possesses a pathway that converts histidine to glutamate (Figure 4; Supplementary Figure 2; Supplementary Table 4). In contrast to most *AqS1* amino acid pathways, the terminal steps of *AqS1* histidine biosynthesis remained highly expressed through the larval phase. Histidinol dehydrogenase (EC 1.1.1.23) was highly expressed in embryos and larvae and declined after settlement, and histidinol-phosphatase (EC 3.1.3.15) peaked in larvae and postlarvae (Figure 4; Supplementary Table 4). On the host side, the terminal enzyme of the histidine degradation pathway, glutamate formimidoyltransferase (EC 2.1.2.5), was significantly upregulated in pre-competent larvae (Supplementary Figure 2; Supplementary Table 4), when phototaxis is maximal (Leys & Degnan, 2001). Two further host–symbiont pairings supported this larval axis. The host’s terminal enzyme is folate-dependent, using tetrahydrofolate as the acceptor for the formimino group released from N-formimino-L-glutamate (Brosnan & Brosnan, 2020), and host and *AqS1* folate metabolism showed reciprocal developmental profiles (Figure 3C,D; Supplementary Table 4). That is, host dihydrofolate reductase (EC 1.5.1.3) was most highly expressed in embryos and lowest in larvae, whereas *AqS1* dihydrofolate reductase was lowest in embryos and peaked in larvae. Also peaking in larvae were *AqS1* 5-formyltetrahydrofolate cyclo-ligase (EC 6.3.3.2), which returns the storage form 5-formyl-THF (folinic acid) to the active folate pool, and several host folate transporters (proton-coupled folate transporters and a mitochondrial folate carrier). Thus, host folate import, symbiont folate output and host demand for folate-dependent histidine catabolism all peaked in the same larval window.

The host also expresses a potential histidine import route, with amino acid and peptide transport being dominated by a markedly expanded family of proton-coupled oligopeptide and histidine transporters, solute carrier family 15 member 4 (SLC15A4), which belongs to the nutrient-exchange transporter set defined below (Supplementary Table 6). Of the 161 SLC15A4-family genes in the *A. queenslandica* genome, 120 were differentially expressed in at least one developmental transition, far more than any other transporter family in this study, and 86 of the 120 (71.7%) had their highest expression in embryos or larvae (Supplementary Table 6, consistent with host import of symbiont-derived amino acids, including histidine, during this period.

### Host and symbiont transporter expression defines two distinct windows of holobiont exchange

To examine the molecular conduits through which metabolites might pass between *A. queenslandica* and its bacterial symbionts, we defined two transporter gene sets in host and symbiont genomes. The first, based on the KEGG PATHWAY ‘membrane transport’ category, comprised 97 host genes, 89% of them ABC transporters and 43 of them multidrug-resistance/xenobiotic-efflux-type transporters (MDR1 and MRP1/3/4/7-family homologues), and 153 *AqS1* genes enriched in sugar, amino acid/peptide and ion transporters; of these, 49 host and 45 *AqS1* genes were differentially expressed across at least one developmental transition. (Figure 5B; Supplementary Figure 3; Supplementary Table 7). Because this set is compositionally dominated by efflux-type ABC transporters in the host, we defined a second, stricter set from the KEGG BRITE ko02000 hierarchy, retaining only families with an established nutrient- or metabolite-transport function and excluding eukaryotic-type multidrug-resistance ABC transporters and other ambiguous categories. This nutrient-exchange set is larger for the host (240 differentially expressed genes) because BRITE, unlike the PATHWAY category, includes the solute carrier (SLC) families, but smaller for *AqS1* (11 differentially expressed genes) (Figure 5A; Supplementary Figure 3; Supplementary Table 6).

**Figure 5.**
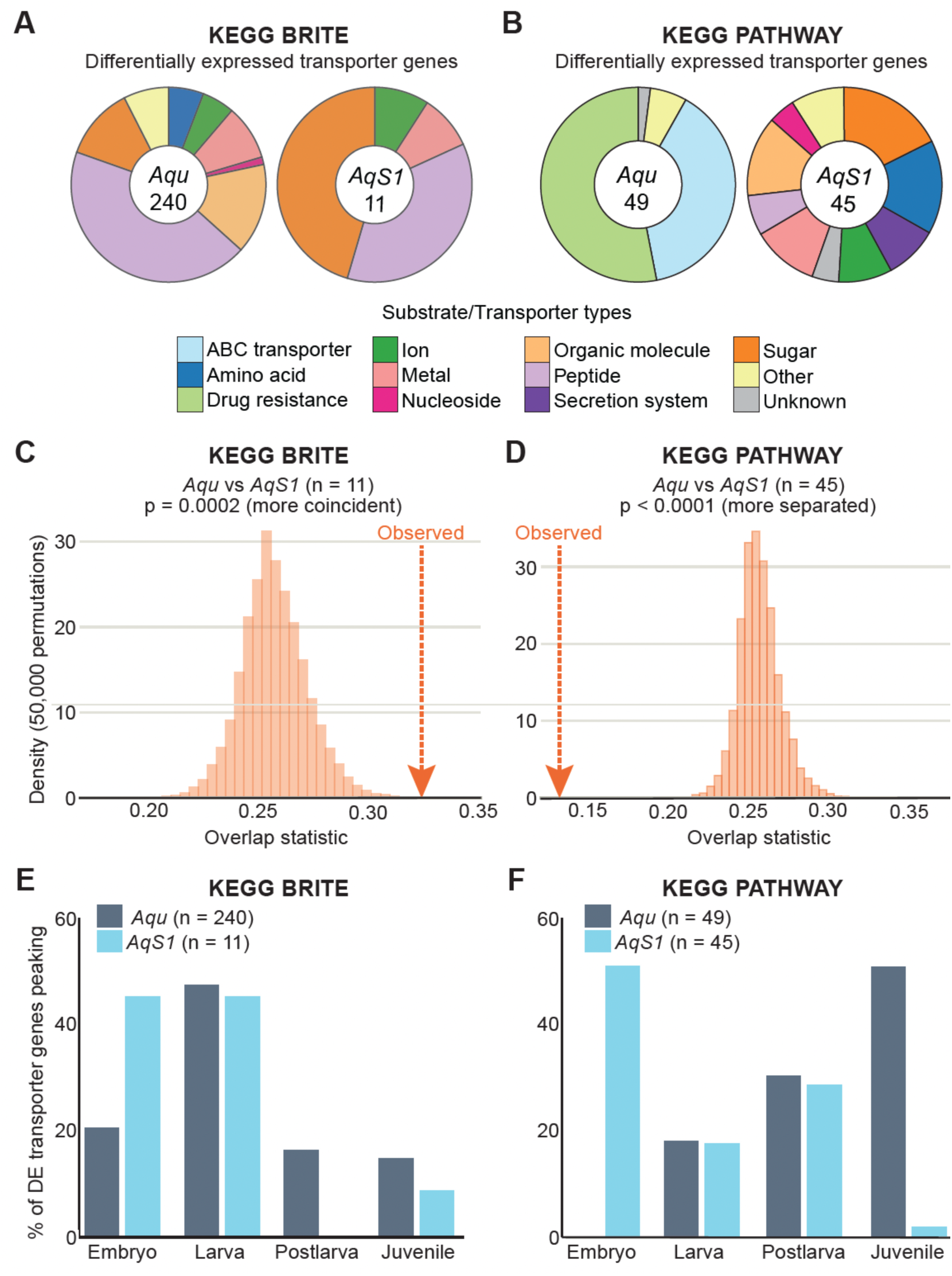
Membrane transporter composition and developmental timing in *A. queenslandica* and its dominant symbiont *AqS1*, compared across two transporter gene sets. The KEGG BRITE set (A, C, E) contains curated nutrient- and metabolite-exchange transporters from ko02000 families with an established transport function. The broader KEGG PATHWAY set (B, D, F) contains the “membrane transport” category. Only genes differentially expressed (|log₂FC| ≥ 1, padj ≤ 0.05) in at least one pairwise developmental comparison are included. (A, B) Composition of each partner’s differentially expressed transporters by substrate or transporter type, with gene counts at centre. (C, D) Overlap in peak-expression stage between *Aqu* host and *AqS1* transporters, compared with a null distribution; orange arrows mark the observed overlap statistic. In the BRITE set, host and *AqS1* transporters peak in the same stages more often than expected by chance (C; one-tailed p = 0.0002). In the PATHWAY set, they peak in different stages more often than expected (D; one-tailed p < 0.0001). (E, F) Percentage of each partner’s differentially expressed transporters peaking at each developmental stage. Note the small *AqS1* BRITE set (n = 11) in E. Volcano plots of differentially expressed transporter genes are shown in Supplementary Figure 3.

For each differentially expressed transporter, we assigned the developmental stage of peak expression and compared host and symbiont peak-stage distributions against a permutation null (50,000 iterations) drawn from each partner’s own whole-transcriptome distribution of peak stages, so that any general skew toward a particular stage was already incorporated in the null expectation. For the nutrient- and metabolite-exchange (KEGG BRITE) set (Supplementary Table 6), host and *AqS1* peak-timing distributions overlapped significantly more than expected by chance (observed = 0.32 versus null mean = 0.26; one-tailed p = 0.0002) (Figure 5C,E). Nutrient and metabolite transporters of the two partners are therefore expressed in coincident developmental windows, as expected if they mediate exchange between them, although the small number of *AqS1* genes in this set (n = 11) limits the resolution of the test.

The broader PATHWAY-based set showed the opposite pattern. *AqS1* differentially expressed transporters peaked overwhelmingly in the embryo (23 of 45, 51%) and almost never in the juvenile (1 of 45, 2%), whereas host differentially expressed transporters peaked predominantly in the juvenile (25 of 49, 51%) and none in the embryo (Figure 5F). *AqS2* reproduced the symbiont pattern with all six of its differentially expressed transporters (Supplementary Figure 3). Under the same permutation framework, host and *AqS1* peak-timing overlapped substantially less than expected by chance (observed = 0.13 versus null mean = 0.26; one-tailed p < 0.0001), and the same held independently for host and *AqS2* (p < 0.0001) (Figure 5D). This separation is driven by the composition of the host set, with the juvenile-peaking host transporters almost entirely MRP/ABCC-family and ABCG efflux-type, which are absent from the nutrient-exchange set (Supplementary Figure 3, Supplementary Table 7). These analyses thus describe different biology, showing the expression of nutrient and metabolite transporters of host and symbiont rise and fall together, whereas the host’s xenobiotic-efflux repertoire and the symbiont’s import repertoire are expressed at host stages that are markedly different developmentally, physiologically and ecologically.

### Host metabolic autonomy increases in the filter-feeding juvenile

The four-day-old juvenile is the only stage sampled that actively filter-feeds, and it is also the stage at which the balance of holobiont metabolic activity shifts most clearly toward the host. The postlarva-to-juvenile transition (i.e., metamorphosis) produced the fewest symbiont differentially expressed genes (63 upregulated, 35 downregulated; Supplementary Figure 1; Supplementary Table 3). This is in contrast to the hundreds of genes that changed at each of the two preceding transitions, indicating that the symbiont transcriptome changes little between 1 hps and the onset of feeding, four days later. The *AqS1* genes that did change were not concentrated in central or amino acid metabolism, with glycan biosynthesis and metabolism being the only KEGG category upregulated in juveniles (Figure 2), suggesting that the symbiont’s response to the onset of host filter-feeding involves remodelling of its cell envelope or extracellular polysaccharides rather than a change in biosynthetic output.

In contrast to the limited change in *AqS1* metabolic gene expression, sponge host central carbon metabolism continued to rise. For instance, glycolytic and oxidative pentose phosphate pathway genes reached their highest expression in juveniles (Figure 3), whereas *AqS1* expression of the same pathways recovered only partially from its larval minimum. The host’s differentially expressed transporter repertoire likewise peaked in the juvenile (25 of 49 genes, 51%; Figure 5, Supplementary Figure 3), driven almost entirely by MRP/ABCC- and ABCG-family efflux-type transporters, while *AqS1* differentially expressed transporters almost never peaked at this stage (1 of 45). Several host and symbiont genes that peaked at settlement declined in juveniles, including host *NOS*, the host zinc efflux transporter SLC30A2, and the terminal enzymes of *AqS1* histidine biosynthesis, which reached their lowest expression in juveniles (Figure 4, Supplementary Figure 2 and 3). Together these patterns describe a juvenile in which host energy metabolism and environmental-interface transport are at their developmental maximum, and symbiont transcription has stabilised.

## Discussion

The three maternally inherited gammaproteobacterial symbionts of *Amphimedon queenslandica*, *AqS1*, *AqS2* and *AqS3* (Fieth et al., 2016), belong to lineages frequently associated with marine sponges (Jia et al., 2026; Kennedy et al., 2008; Kennedy et al., 2014; Schmitt et al., 2012; Taylor et al., 2007; Thomas et al., 2016). Together these symbionts contribute almost half of the hologenome’s metabolic genes despite encoding only 20.4% of its total KEGG-assigned genes (Xiang et al., 2026).

This disproportionate contribution underscores the functional dependence of this animal on its vertically-inherited microbiome, with symbionts completing host amino acid, cofactor/vitamin, glycan, carbohydrate, lipid and inorganic nutrient metabolic pathways. In the adult sponge, these pathways are transcriptionally active and are complemented by the expression of membrane transporters predominantly encoded by symbionts, supporting an exchange-based architecture for nutrient cycling across the holobiont (Xiang et al., 2026). Shifts in community composition are known to occur during the *A. queenslandica* life cycle (Fieth et al., 2016), and may be one way in which the microbiome directly affects holobiont function (Petrullo et al., 2026). Here, we focus on the symbiont’s side of the partnership to identify a second way in which core symbionts may contribute to holobiont function throughout the life cycle, via changes in their transcriptional activity.

The transcriptomes of each of the four holobiont partners that we assayed were strikingly stage-specific across embryonic, larval and metamorphic development, suggesting that symbiont gene expression is closely coupled to host ontogenetic state (Figure 6). It is unclear if this influence is via specific host-derived signals or is a response to overall changes in the holobiont’s biochemical environment. Although the magnitude of the partners’ contributions cannot be inferred from these normalised expression data, this analysis shows that symbiont biosynthetic gene expression tracks with host developmental stage, and may extend beyond nutrition to the developmental transitions themselves.

**Figure 6.**
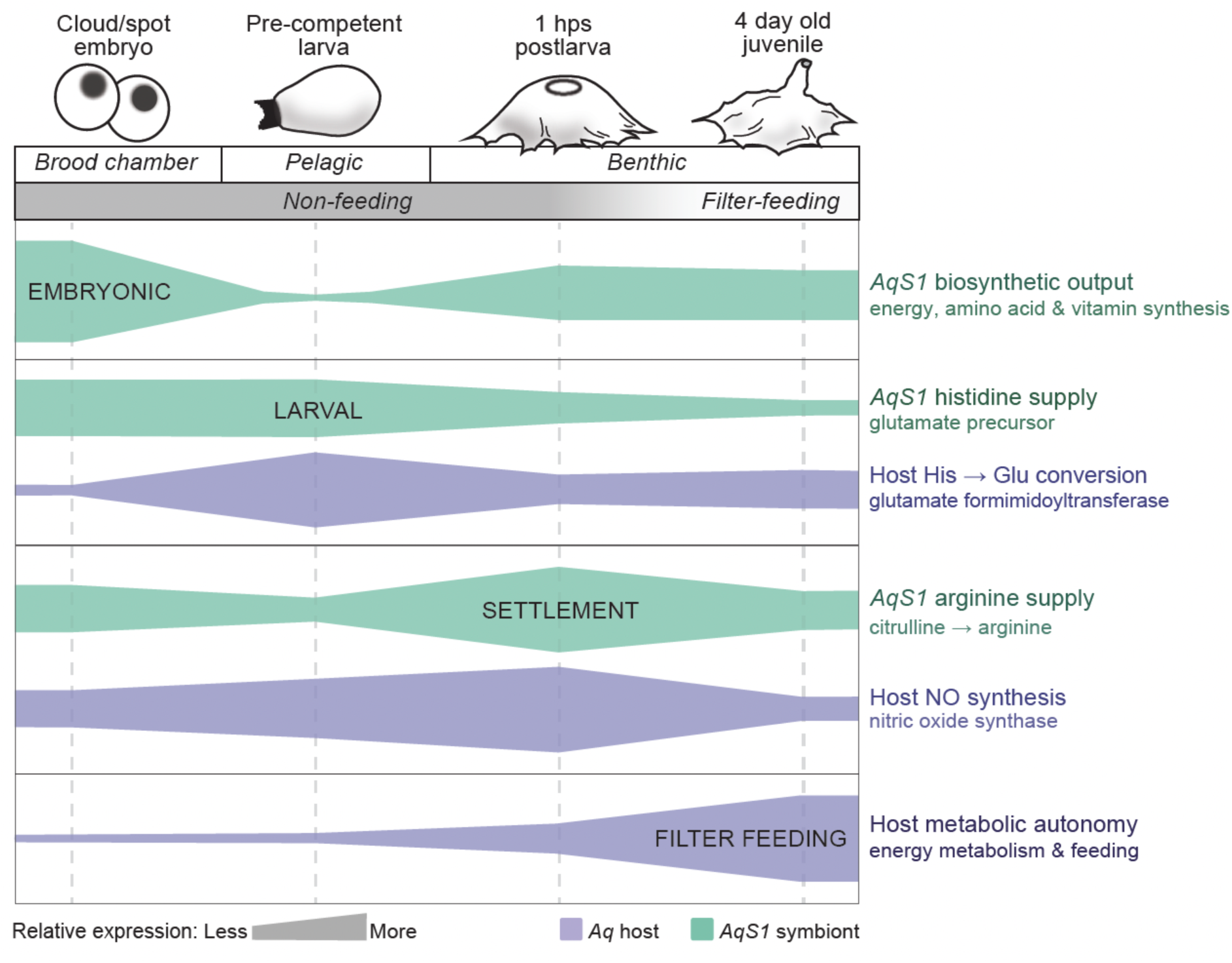
Summary: symbiont biosynthetic output is scheduled to host developmental stage. Band width shows relative expression within each partner across the four developmental stages (*AqS1*, green; *A. queenslandica*, purple). Bands are schematic summaries, and widths are not comparable between partners. *AqS1* biosynthetic output peaks in the brooded embryo, contracts in the non-feeding larva and partially recovers after settlement. The larval contraction is selective. Symbiont histidine supply is maintained in the larva, coinciding with host histidine-to-glutamate conversion. Symbiont arginine supply is held low in the larva, when *AqS1* nitric oxide reductase also peaks, and rises at settlement together with host nitric oxide synthase, consistent with a precursor for the nitric oxide signal that initiates larval settlement. Host metabolic autonomy increases through metamorphosis to a maximum in the filter-feeding juvenile.

### Symbiont biosynthetic output peaks during embryogenesis

The high fidelity vertical transmission of the core symbionts during oogenesis and early embryogenesis means they are present from the earliest stage of development (Fieth et al., 2016). Embryos develop in brood chambers, nourished by maternal yolk rich in lipids and proteins (Koutsouveli et al., 2022), while undergoing energetically demanding cell division and differentiation, and morphogenesis (Adamska et al., 2007; Degnan et al., 2008). While host energy-metabolism genes generally are highly expressed throughout development, *AqS1* glycolytic and TCA cycle genes are at their highest expression in embryos, consistent with ready access to maternal yolk-derived carbon sources, and suggesting that the symbiont’s own energy metabolism is most active at this stage. This is in contrast to the host’s, which is generally highest in the filter-feeding juvenile. The PPP pathway comprises an oxidative branch that generates NADPH, and a non-oxidative branch that generates nucleotide precursors and reversibly interconverts sugar phosphates and feeds them into glycolysis (Stincone et al., 2015). Of the symbionts, only *AqS1* encodes and expresses the complete oxidative branch, and during embryogenesis, when host oxidative PPP expression is at its lowest, *AqS1* oxidative PPP expression peaks. This reciprocal timing raises the possibility that symbiont-derived NADPH, or the products of NADPH-dependent biosynthesis, supplement the host at this stage.

*AqS1*’s nutritional contribution during embryogenesis appears to take two forms. The first is biosynthetic, with peak expression of *AqS1* amino acid, vitamin and central-carbon genes, and potentially associated transporters. The second is direct consumption, since *A. queenslandica* embryonic cells phagocytose *AqS1* (Fieth et al., 2016), as sponge cells do symbionts in other species (Díez-Vives et al., 2022; Riesgo et al., 2007; Turon et al., 2025). Recent work in *Halichondria panicea* shows how far the second mode can go, with nurse cells phagocytosing the dominant symbiont and likely converting it to yolk, so that the symbiont’s abundance falls (Turon et al., 2026). In *A. queenslandica, AqS1* abundance is maintained through embryogenesis and into the larva (Fieth et al., 2016), and genes peaking in the embryo are enriched for enzymatic functions, consistent with the symbiont being primarily a metabolic producer at this stage and only secondarily a food source. The two sponges may thus illustrate two ends of a continuum in how a maternally provisioned embryo draws on its microbiome, from harvesting symbiont-secreted metabolites to harvesting the symbionts themselves.

In cloud/spot embryos, peak *AqS1* expression of amino acid biosynthetic genes, including those for histidine, coincides with expression of host SLC15A4-family transporters, which are best characterised for histidine and peptide translocation (Custódio et al., 2023; Zhang et al., 2025). Most of the differentially expressed members of this expanded family peak in embryos or larvae, which suggests that the host can import symbiont-derived amino acids from embryogenesis onwards. Folate (vitamin B₉) metabolism shows a similar complementarity, but with a hand-over between stages. The host encodes only part of the pathway and expresses its final step, dihydrofolate reductase, most highly in embryos. In larvae, *AqS1* folate biosynthesis genes dominate, together with increased expression of host folate transporters. This reciprocal pattern would be consistent with continuity of folate supply into the non-feeding larval stage, and links directly to the larval histidine–glutamate axis discussed below. Together, these pairings suggest that stage-specific expression of symbiont biosynthetic genes is matched by host capacity to take up their products.

Similar to the complementation for folate biosynthesis, *A. queenslandica* encodes the terminal synthesis steps of multiple amino acid pathways — phenylalanine, tyrosine, methionine, isoleucine, leucine, valine and lysine — despite lacking the upstream intermediary enzymes. The partitioning may be enhanced further by having paralogue redundancy at the final enzymatic step (Xiang et al., 2026), potentially allowing the host to modulate amino acid availability in response to endogenous developmental or environmental signals.

### The larval holobiont enters a state of shared metabolic constraint

*A. queenslandica* larvae emerge from brood chambers in a pre-competent state and spend several hours in the plankton before becoming competent to settle (Degnan & Degnan, 2010). The emergence into the water column coincides with a strong shift in symbiont gene expression, with *AqS1* downregulating genes involved in glycolysis, TCA cycle, amino acid biosynthesis, pantothenate and CoA synthesis, and ATP generation via the AprAB/Sat module of its sulfur oxidation pathway (Figure 8). This broad downregulation of symbiont biosynthetic gene expression is unlikely to reflect a decline in symbiont abundance because *AqS1* abundance is maintained in larvae despite some phagocytosis by sponge cells (Fieth et al., 2016). Thus the reduction appears to reflect a change in the physiological state of the symbionts.

Sponge larvae are non-feeding, such that metabolic contraction of *AqS1* in this phase may reflect energy conservation, reducing energetically-expensive biosynthesis in a resource-limited stage of development; larval yolk reserves are needed for larval swimming and metamorphosis (Degnan et al., 2008). Consistent with this interpretation, *AqS1* in larvae upregulates its import machinery, including multiple sugar, amino acid and peptide transporters, as well as C4-dicarboxylate transporters that facilitate the uptake of TCA intermediates like succinate, malate and fumarate (Janausch et al., 2002). This suggests that as the symbiont shifts away from biosynthesis during the larval phase, it becomes increasingly dependent on host-derived substrates. Thus the contraction appears to be a holobiont-wide response to a shared resource limitation rather than a symbiont strategy per se. As described below, however, this contraction is not universal; several *AqS1* biosynthetic pathways track with host developmental requirements, consistent with pathway-specific gene regulation operating in *AqS1* despite the general energy-conservation mode. The same selectivity extends to *AqS1* sulfur energy metabolism. Constitutive *sox* expression indicates that the symbiont does not shut down in the larva, whereas repression of the ATP-generating AprAB/Sat module parallels its biosynthetic contraction; the significance of the concurrent larval peak in *dsr* expression remains to be resolved. The transcriptional switch at settlement is both the release of the whole holobiont from a resource-limited state, and the induction of specific symbiont pathways whose products are required by the host.

### Symbiont arginine biosynthesis is timed to settlement

Metamorphic competence in marine invertebrates is characterised by the activation of chemosensory receptors, signalling pathways and transcriptional regulators that detect and respond to benthic settlement cues (Degnan & Degnan, 2010; Jackson et al., 2002; Jensen & Morse, 1990; Morse et al., 1980; Yang et al., 2026). In *A. queenslandica*, G-protein coupled receptors (GPCRs) and associated signal transducer genes, along with regulatory transcription factors, are activated as competence develops about 6 hours after larvae emerge from the maternal adult (Say & Degnan, 2020; Yuan et al., 2026). Nitric oxide (NO) and MAPK signalling are necessary for the induction of settlement and metamorphosis (Ueda et al., 2016). Prior to the acquisition of competence, pre-competent larvae are maximally negatively phototactic (Leys and Degnan, 2001). In the current study, our data extend the evidence for a symbiont role in settlement and suggest a candidate second link, to larval glutamate signalling.

Song et al. (2021) demonstrated experimentally that arginine biosynthesis by *A. queenslandica* symbionts is required for NO production and larval settlement. In the present study, we show that *AqS1* expression of the terminal arginine biosynthesis enzyme (argininosuccinate lyase) lacks the embryonic peak seen in other pathways, and increases instead around the time of settlement (i.e. by 1 hps postlarvae) when host NOS expression is also peaking (Figure 8). This citrulline-to-arginine two-step process cannot be performed by the host and is uniquely upregulated in *AqS1* in time for settlement, unlike earlier steps in this biosynthetic pathway, which follow the high embryonic expression pattern observed for most metabolic pathways. Because host NOS itself generates citrulline as it produces NO, this symbiont module is positioned to regenerate arginine from the host’s own NO-derived citrulline, analogous to the citrulline–NO cycle by which vertebrate cells sustain high-output NO synthesis through coordinated expression of NOS, argininosuccinate synthase and argininosuccinate lyase (Husson et al., 2003). Our previous work (Song et al., 2021) provided empirical support for holobiont complementation of the arginine-citrulline loop, and the settlement-timed gene activation reported in the current study is consistent with this. We are unaware of other examples of the citrulline–arginine recycling being split between an animal host and a bacterial symbiont, but the aphid–*Buchnera* arginine pathway is an analogous symbiosis precedent for host–symbiont partitioning of a single pathway with the host retaining one step (Hansen & Moran, 2011).

Although the suppression of arginine synthesis in pre-competent larvae might simply be following the general pattern of biosynthetic repression at this stage (described above), two observations argue against this. First, the citrulline-to-arginine module does not peak in the embryos as most other *AqS1* biosynthetic pathways do, and, second, *AqS1* nitric oxide reductase, which degrades NO, peaks in the larva, in contrast to the general trend of declining expression at this stage. The two *AqS1* gene sets with the most direct bearing on NO availability therefore both depart from the bulk developmental expression pattern, one which would suppress NO in the pre-competent larva and the other which makes the necessary NO precursor available in competent larvae. Together, these expression profiles lend further support to our proposal that symbiont arginine biosynthesis is a point of symbiont-mediated regulation of a host developmental transition (Song et al., 2021). Whether this timing is imposed by host-derived signals or by cues from the postlarval environment cannot be resolved from expression data, although diel regulation of symbiont arginine biosynthesis in the demosponge *Aplysina aerophoba* (Ramírez et al., 2026) shows that environmental control of this pathway is plausible. We propose a testable model in which the combined effect of restricted arginine supply and active NO degradation could result in NO concentration being maintained in pre-competent larvae below the threshold required for the induction of settlement and metamorphosis, allowing the larva to further disperse in the water column. In this scenario, *AqS1* arginine biosynthesis resumes in competent larvae approaching settlement, NOR expression declines, and the NOS-NO axis in the host tips into the range necessary for settlement and metamorphosis.

A second, as yet untested, link concerns histidine. *AqS1* histidine biosynthesis genes are maintained in pre-competent larvae, when the host’s folate-dependent histidine-to-glutamate conversion step is upregulated and *AqS1* folate biosynthesis and host folate import also peak. The symbiont therefore has the capacity to contribute both the substrate and the co-enzyme for a host glutamate pathway implicated in both larval phototaxis and settlement (Say and Degnan, 2020; Wong et al., 2022; Zhang et al., 2026).

### Transporter expression defines two distinct windows of holobiont exchange

Transporter expression provides transcriptional evidence that the metabolic complementarity described above is accompanied by the machinery to move metabolites between partners. When the analysis is restricted to transporter families with an established nutrient- or metabolite-exchange function, host and *AqS1* transporters peak in coincident developmental periods significantly more often than expected by chance. Although coincident expression does not by itself demonstrate exchange, this is the pattern predicted if the two partners’ transport programmes are co-ordinated around exchange. It is consistent with the specific pairings identified as case studies, including (i) host SLC15A4-type histidine/peptide transporters peaking in embryos and larvae when *AqS1* amino acid biosynthesis is high, (ii) host folate transporters and *AqS1* folate biosynthesis both peaking in larvae; and (iii) a bacterial zinc importer and host zinc exporter peaking together at settlement, consistent with host-to-symbiont zinc transfer at that stage. *AqS1* C4-dicarboxylate transporters, upregulated in larvae, similarly point to the symbiont drawing on host TCA intermediates during the period of its own biosynthetic restriction.

A second, quite different pattern emerges from the broader transporter set, in which symbiont and host transporters occupy distinct stages of the life cycle: *AqS1* and *AqS2* upregulate their transporter repertoires predominantly in embryogenesis, coincident with the period when these symbionts are biosynthetically most active and import capacity for substrate acquisition would be at a premium, whereas the host’s transporter repertoire expands sharply in the juvenile. This difference is driven almost entirely by host MRP/ABCC- and ABCG-family transporters, whose best-characterised roles are in xenobiotic and metabolite efflux rather than nutrient uptake. Their induction in the juvenile coincides with the onset of filter-feeding, when the sponge first pumps seawater through its body and is exposed to environmental microbes, their metabolites and dissolved xenobiotics. Consistent with these results, experimental introduction of foreign bacteria to juvenile *A. queenslandica* induces a strong xenobiotic response, which is in stark contrast to the innate immune response induced by native symbionts (Yang et al., 2026). We therefore interpret the juvenile expansion of host efflux transporters not as a change in host–symbiont exchange, but as the onset of an environmental-defence programme that accompanies feeding. This is markedly different from the host’s response to vertically-inherited constitutive symbiont community established during oogenesis and early embryogenesis (Fieth et al. 2016; Yang et al., 2026). Thus the nutrient- and metabolite-exchange machinery of holobiont partners shows coincident peaks across development more often than expected by chance, while a separate host efflux system switches on when the animal opens itself to the environment.

### Filter-feeding reduces symbiont metabolic dependency

Metamorphosis in sponges culminates in the formation of the aquiferous system, enabling juveniles to filter-feed directly from seawater (Bergquist & Green, 1977; Degnan et al., 2015; Sogabe et al., 2016). The postlarva-to-juvenile transition occurs without a change in environment and has the fewest symbiont DEGs of any stage transition in this study. This suggests that it is the pelagic-to-benthic ecological transition, rather than ontogenetic progression per se, that most powerfully reshapes the symbiont transcriptome. At the onset of feeding, *AqS1* more strongly expressed glycan biosynthesis and metabolism genes compared to other stages. This may reflect a remodelling of the symbiont’s cell envelope or extracellular polysaccharides as the sponge is exposed for the first time to a continuous flow of environmental bacteria, and to the host’s expanding phagocytic and efflux machinery, both of which rely on sponge cells discriminating amongst bacteria on the basis of their surface properties (Pita et al., 2018; Wehrl et al., 2007). This is consistent with the ability of filter-feeding juvenile *A. queenslandica* to differentially process native and foreign bacteria (Yang et al., 2026). Whether this would represent a competitive, protective or recognition-related response cannot be resolved from the current expression data, but it indicates that the juvenile transition, while metabolically quiet for the symbiont, is not without a symbiont-side response. The juvenile is the only stage in which host metabolic activity and transport capacity exceed those of the symbiont – host glycolytic and PPP genes are upregulated, consistent with carbohydrate metabolism being fuelled by ingested material from seawater. The oxidative PPP steps, expressed only by *AqS1* during embryogenesis when the host’s own oxidative branch is less active, are now fully active in the host itself.

### An ancient developmental symbiosis?

Nitric oxide signalling has been proposed as an ancient regulator of the animal pelagobenthic life cycle, on the basis of its role in settlement across sponges, cnidarians and bilaterians (Ueda et al., 2016). The present study identifies a vertically inherited bacterial symbiont as a likely source of the NO precursor (Song et al., 2021), and shows that this supply tracks to a specific host developmental stage. Given the phylogenetic position of sponges and the probable antiquity of the biphasic life cycle (Boyle & Young, 2025; Degnan & Degnan, 2023), one reading is that symbiont-supplied precursors for developmental signalling were a feature of the earliest animal holobionts. The alternative, that such dependencies have arisen independently in sponges, in the squid–*Vibrio* light-organ symbiosis and in the mammalian gut, cannot be excluded on present evidence; the symbiont lineages involved are unrelated, and the pathways recruited differ. Distinguishing these possibilities will require comparable partner-resolved developmental transcriptomes from other early-branching lineages, and from other sponges with vertically-transmitted symbionts; the recent demonstration of symbiont phagocytosis supporting embryogenesis in *H. panicea* (Turon et al., 2026) already indicates that microbial integration into early development is not peculiar to *A. queenslandica* (Fieth et al., 2016). Importantly, this study suggests that a microbiome can influence successive stages of an animal life history (Petrullo et al., 2026) not by changing in composition but by reprogramming its transcriptional activity. That this occurs in a sponge, with an ancestral-type biphasic life cycle and without the specialised host systems that mediate microbial influence on development in bilaterians, is consistent with developmental symbiosis being an early and possibly ancestral feature of the animal holobiont.

## Methods

### Sample collection and RNA extraction

Adult *A. queenslandica* were collected from Heron Reef, Australia (23.44°S, 151.92°E) and maintained in a closed aquarium at the University of Queensland under GBRMPA permit G16/38120.1; the four developmental stages were procured as previously described (Leys et al., 2008; Say & Degnan, 2020; Sogabe et al., 2016). Cloud and spot stage embryos (n=60) were collected from dissected adult brood chambers. Swimming pre-competent larvae (n=60) were collected within 1 hour post-emergence (1 hpe) from adult sponges. Postlarvae settled and metamorphosing on the articulated coralline alga *Amphiroa fragilissim*a were sampled at 1 hour post settlement (1 hps; n=60). Juveniles, identified by the presence of an osculum, were sampled at 96 hps (n=60). All samples were stored in RNA Later (Invitrogen) and RNA was isolated from three pools of 20 individuals for each developmental stage using Trizol® reagent (ThermoFisher) according to the manufacturer’s protocols with slight modification: 1-Bromo-3-chloropropane (Sigma Aldrich) was used as the phase separation agent instead of chloroform.

### Library preparation and sequencing

To enrich for the microbial RNA from these 12 pooled biological replicates from four developmental stages (n=3 replicates of 20 pooled individuals per stage), poly(A) RNA was depleted using the Poly(A) Purist^TM^ MAG kit (ThermoFisher) for 30 minutes and the resultant poly(A)-depleted RNA was purified with SPRI Magnetic Beads (RNAClean XP, Beckman Coulter). rRNA from the poly(A)-depleted RNA was removed using customised riboPOOL (siTOOLS). cDNA libraries were constructed from the final poly(A)-depleted, rRNA-depleted RNA using the TruSeq stranded total RNA kit (Illumina) and unique dual indexes (UDI; Illumina), quantified and quality checked with a BioAnalyzer (Agilent), and sequenced with NovaSeq (Genome Innovation Hub, University of Queensland).

### Read processing, mapping and counting

The quality of raw RNA sequencing reads from the developmental libraries were checked using FastQC (Andrew, 2010) and sequences were trimmed with Trimmomatic (v0.36) (Bolger et al., 2014). Trimmed reads were mapped to improved host and symbiont genomes (Xiang et al., 2026) — the *A. queenslandica* genome (GCA_016292275.1) using HISAT2 (v2.0.5) (Kim et al., 2019), and *AqS1* (GCA_015278165.1), *AqS2* (GCA_015278175.1) and *AqS3* (GCA_015278115.1) using Bowtie2 (Langmead & Salzberg, 2012). Reads were counted with HTSeq (v1.0) (Anders et al., 2014) for *A. queenslandica* and RSubread (v1.22.2) featureCounts (Liao et al., 2014) for the symbionts.

Normalisation of raw count data was achieved using the DESeq2 R package (v1.44.0) (Love et al., 2014), filtering out genes with fewer than 5 counts across all samples. Normalisation was performed independently for each holobiont partner, so that expression comparisons across developmental stages within a partner are not affected by changes in the relative read contribution of the other partners.

### Principal component analysis and differential expression analysis

A regularised log transformation (rlog) was applied to the normalised counts and distance matrices were computed using Euclidean metrics. Principal component analysis (PCA) was performed on the top 500 most variable genes using the *prcomp* function in R and visualised using the factoextra R package (v2.0.0) (Kassambara & Mundt, 2026).

Differentially expressed genes (DEGs) were identified from normalised count data using the DESeq2 R package (v1.44.0) (Love et al., 2014). A gene was considered differentially expressed if it met the criteria of |log_2_FC| ≥ 1 and adjusted p-value (padj) ≤ 0.05. Volcano plots of DEGs were generated using the EnhancedVolcano R package (v1.22.0) (Blighe et al., 2024).

### Functional annotation and pathway mapping

Unless otherwise stated, all KEGG Orthology (KO) assignments and Enzyme Commission [EC] numbers were taken from Xiang et al. (2026). KO-annotated genes were mapped onto KEGG pathways using KEGG Mapper Reconstruct (Kanehisa & Sato, 2020) and by screening against the KEGG Orthology hierarchy file (ko00001.keg; release 119.0 (Kanehisa et al., 2023). Where multiple genes from one genome were assigned a KO number corresponding to the same EC number, the gene with the highest mean normalised expression count was used as a representative for pathway activity.

### Expression profiling

Genes were ranked based on the average normalised gene expression values across replicates. The dataset was divided into four equal groups, ranging from the lowest quartile (Q1), containing lowly- and non-expressed genes, to the highest quartile (Q4), containing the top 25% most highly expressed genes. Due to low gene coverage in *AqS2* and *AqS3* datasets, only genes expressed in Q3 or Q4 in at least one developmental stage were included in further analysis.

Gene expression heat maps were generated using the Pheatmap R package (v1.0.12) (Kolde, 2018). Z-scores were computed per gene, across the four stage means, separately within each partner.

### Peak-stage assignment

For each gene, the developmental stage (embryo, larva, postlarva or juvenile) at which mean normalised expression across replicates was highest was designated its peak stage. Peak stages were assigned separately for each holobiont partner, to all expressed genes, and the resulting whole-transcriptome distribution of peak stages for each partner was used as the background for the analyses below.

### Enzyme-annotation peak-stage enrichment

To test whether the embryonic peak in *AqS1* metabolic gene expression reflected a general elevation in transcriptional activity at this stage or a specific enrichment of enzymatic function, *AqS1* genes were classified as enzyme-encoding (assigned an EC number; Supplementary Table 5) or non-enzymatic (no EC). Because the peak stage of lowly expressed genes is assigned less reliably, the analysis was restricted to genes in the upper two expression quartiles (Q3 or Q4) in at least one developmental stage (n = 1,858; 787 enzyme-encoding, 1,071 non-enzymatic). Each gene was assigned to the stage of highest mean normalised expression (as above), and independence between enzyme annotation and peak stage was tested with a chi-square test on the 2 × 4 contingency table. Stage-specific contributions to the overall association were assessed from adjusted standardised residuals: a residual > 1.96 indicates a stage-specific departure at α = 0.05. For the embryo, the odds ratio for enzyme-encoding versus non-enzymatic genes peaking at that stage rather than at any other was calculated, with a 95% Wald confidence interval. Because genes co-transcribed within operons are not independent, p-values should be regarded as indicative rather than exact.

### Transporter gene sets and permutation analysis

Two transporter gene sets were defined for the host and each symbiont. The first comprised genes assigned to the KEGG PATHWAY ‘membrane transport’ category (ABC transporters, map02010; phosphotransferase system, map02060; bacterial secretion system, map03070) (Supplementary Table 7). The second, stricter set was curated from KEGG BRITE ko02000 families with an established nutrient- or metabolite-transport function (selected SLC, MFS, prokaryotic-type ABC, PTS and primary active transporter [TC:3] subfamilies), excluding eukaryotic-type multidrug-resistance-associated ABC transporters and other ambiguous categories (Supplementary Table 6). In each set, genes meeting the differential expression criteria above in at least one pairwise developmental comparison were retained.

Overlap in peak-timing between the host and a given symbiont was summarised as Σᵢ(aᵢ⁄n_a)(hᵢ⁄n_h), where aᵢ and hᵢ are the numbers of symbiont and host genes peaking in stage i and n_a and n_h are the corresponding set sizes. A null distribution was generated by 50,000 permutations in which symbiont and host gene sets of the observed sizes were redrawn by multinomial sampling from each partner’s whole-transcriptome peak-stage distribution, so that any partner-specific baseline skew toward a particular stage was incorporated in the null expectation. One-tailed p-values were the proportion of permutations with a statistic at or below the observed value (testing for separation) or at or above it (testing for coincidence). The analysis was run for host–*AqS1* and host–*AqS2*; *AqS3* yielded a single differentially expressed transporter and was excluded. Analyses were performed in Python [version 3.12.7] with pandas [version 2.2.3] and NumPy [version 2.2.2]; code is available at Zenodo [DOI: 10.5281/zenodo.22934393]. Four transporter families with direct links to biosynthetic pathways examined in this study (SLC15A4-type peptide/histidine transporters, zinc uptake and efflux, folate transport and C4-dicarboxylate transport) were additionally examined individually using the same differential expression criteria.

## Supporting information

List of Supplementary Tables

## Declarations

## Ethics approval and consent to participate

Not applicable

## Consent for publication

Not applicable

## Availability of data and materials

The dual RNASeq raw read datasets generated for the current study are available at NCBI under Bioproject PRJNA982967. All read count data that supports the findings of thi study are included in the Supplementary Tables associated with this article.

## Authors’ contributions

Eve Maunders: methodology, visualisation, Writing – original draft preparation; Sheen Wong: generation of dual RNASeq data, methodology, visualisation; Rebecca Fieth: electron microscopy; Xueyan Xiang: genome assemblies and functional annotations for all holobiont partners; Bernard M Degnan: conceptualisation, methodology, funding acquisition, resources, project administration, supervision, Writing – original and final draft preparation; Sandie M Degnan: conceptualisation, methodology, funding acquisition, resources, project administration, supervision, Writing – original and final draft preparation.

## Acknowledgements

The authors thank staff at the Heron Island Research Station for logistical assistance during field collections, and Nick Rhodes at QCIF for management of high-performance computing facilities.

