## Supplementary Figures for "Symbiont biosynthetic output tracks host developmental stage in a marine sponge holobiont"

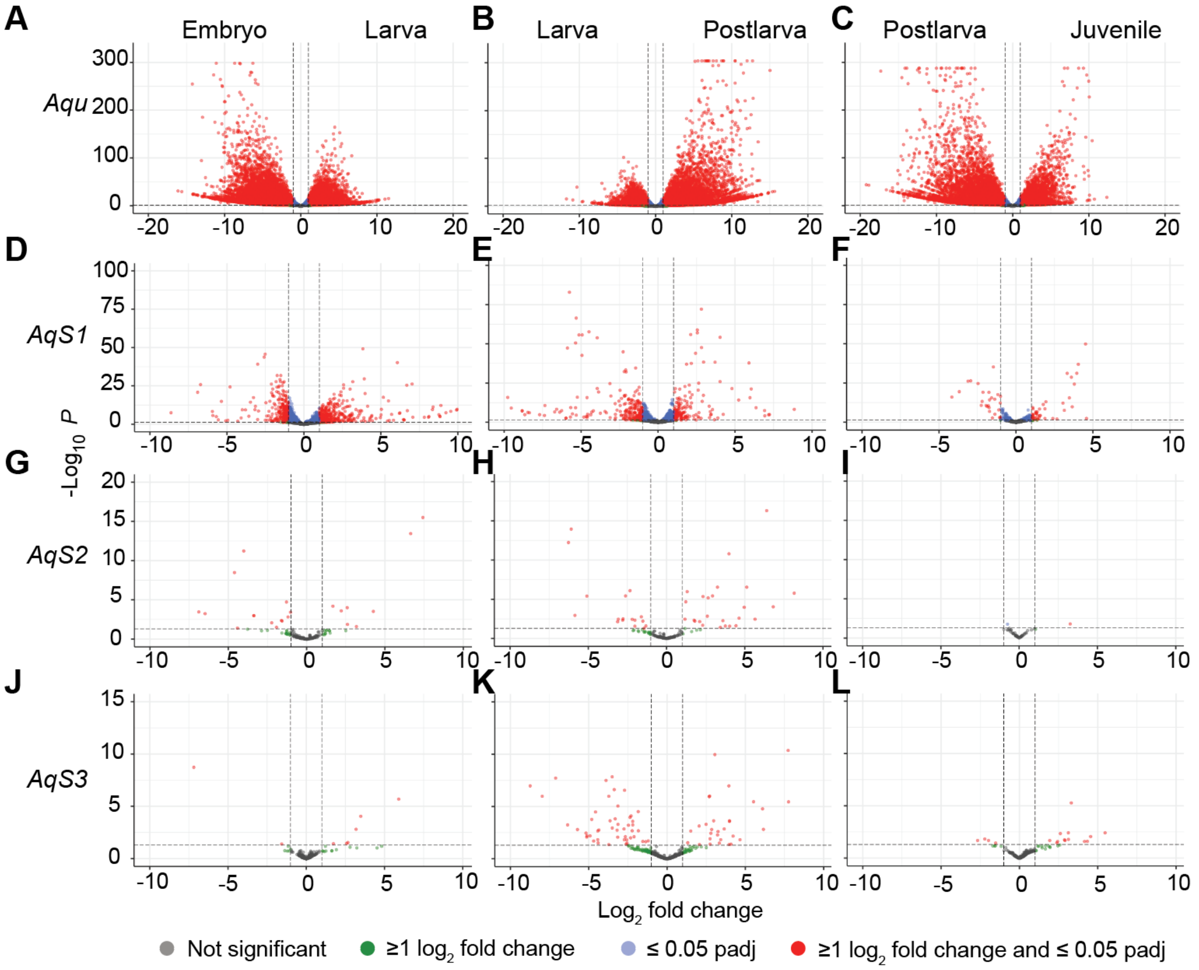


**Supplementary Figure 1.** Volcano plots of differentially expressed genes in host and symbionts across embryonic, larval, 1 hps postlarval and juvenile developmental stages. **(A-C)** *A. queenslandica* (*Aqu*); **(D-F)** *AqS1*; **(G-H)** *AqS2*; and **(J-L)** *AqS3*. Significantly differentially expressed genes (|$\log_2$FC|≥1 and padj ≤ 0.05) are in red. Only genes expressed in the top two quartiles (Q3 or Q4) in at least one developmental stage are included in *AqS2* and *AqS3* analysis, to account for low sequencing depth for those symbionts. **(A, D, G, J)** Differentially expressed genes between cloud/spot embryonic (left) and pre-competent larval (right) stages, with genes upregulated in embryos and larvae to the left and right of the centre line, respectively. **(B, E, H, K)** Differentially expressed genes between pre-competent larval (left) and 1 hps postlarval (right) stages. **(C, F, I, L)** Differentially expressed genes between 1 hps postlarval (left) and juvenile (right) stages.

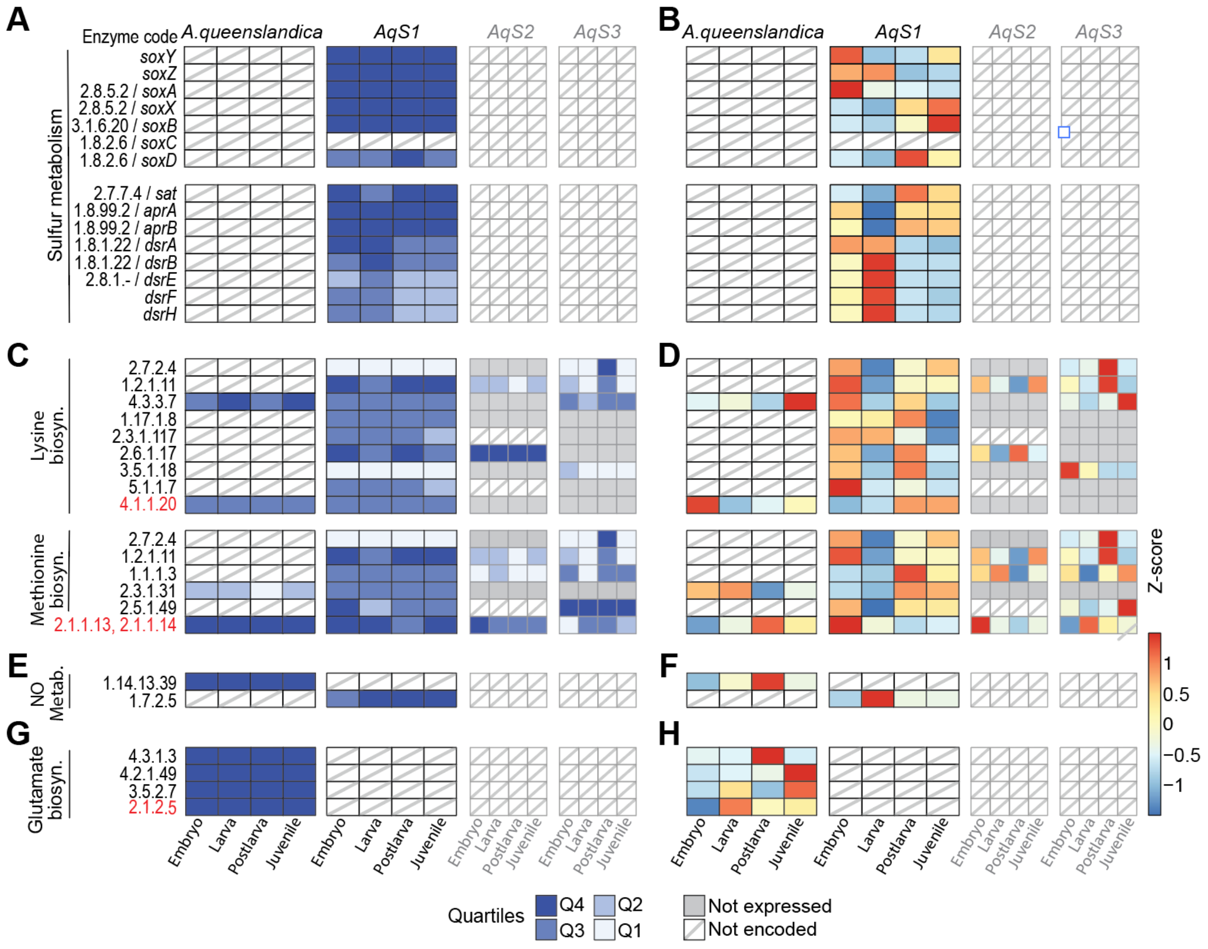


**Supplementary Figure 2.** Expression of sulfur metabolism, lysine, methionine and glutamate biosynthesis, and nitric oxide metabolism genes in *A. queenslandica* and its bacterial symbionts (*AqS1*, *AqS2* and *AqS3*) across embryonic, larval, 1 hps postlarval and juvenile developmental stages. **(A, B)** Sulfur metabolism; **(C, D)** lysine and methionine amino acid biosynthesis; **(E, F)** nitric oxide (NO) metabolism; and **(G, H)** glutamate biosynthesis. (**A, C, E, G)** Expression levels are shown as quartiles, with Q4 representing the top 25% of expressed genes within each transcriptome. **(B, D, F, H)** Z-scores normalised individually for each holobiont partner, per gene. Enzyme codes coloured in red represent the final enzyme in the biosynthesis pathway.

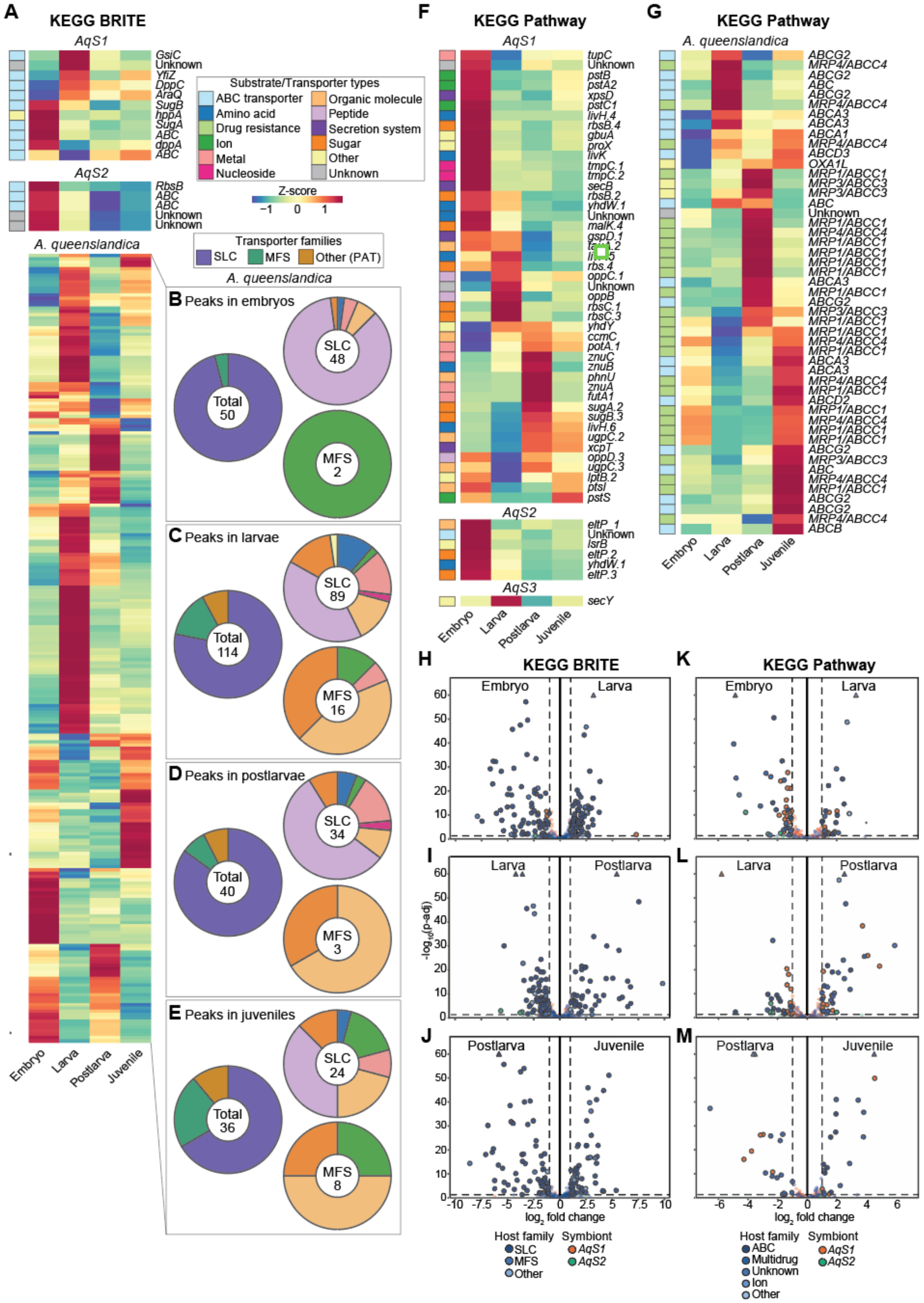

**Supplementary Figure 3.** Analysis of significantly differentially expressed transporter genes in *A. queenslandica* and its symbionts (*AqS1, AqS2* and *AqS3*) across embryonic, larval, 1 hps postlarval and juvenile developmental stages ($|\log_2FC|\geq1$ and padj $\leq 0.05$). **(A-E)** Analysis of differentially expressed nutrient- and metabolic-transporter genes identified by KEGG BRITE. **(A)** Heat map of differentially expressed genes in *AqS1*, *AqS2* and *A. queenslandica*. The coloured bar to the left of the *AqS1* and *AqS2* heat maps ascribes each gene to a broad substrate/transporter family that is listed in the key to the right of the heat maps. For *A. queenslandica*, differentially expressed transporter genes are parsed into four groups based on the developmental stage at which their expression is maximal/peaks (B-E). **(B)** Genes with peak expression in cloud/spot embryos. The donut chart to the left parses the total number of genes (50) into either SLC transporters (purple, 48), MFS transporters (green, 2) or Other (PAT) transporters (brown, 0). The donut chart in the upper right parses the 48 SLC transporters into broad substrate/transporter families (the colour corresponds to the key above). The chart in lower right parses the MLFs into families. **(C)** Donut charts of genes with peak expression in pre-competent larvae; **(D)** genes with peak expression in 1 hps postlarvae; **(E)** genes with peak expression in juveniles. **(F, G)** Heat maps of differentially expressed efflux, xenobiotic and other transporter genes identified by KEGG PATHWAYS, with all genes named to the right of the heat map. **(F)** Symbiont genes; **(G)** *A. queenslandica* genes. The coloured bar to the left of the heat maps ascribes each gene to a broad substrate/transporter family that is listed in the key to the left of the heat maps. **(H-M)** Volcano plots of differentially expressed transporter genes in host and symbionts across embryonic, larval and metamorphic development. **(H, I, J)** Transporters identified by KEGG BRITE. **(K, L, M)** Transporters identified by KEGG PATHWAYS. **(H, K)** Pairwise comparison of differential gene expression between cloud/spot embryonic (left) and pre-competent larval (right) stages, with genes upregulated in embryos and larvae to the left and right of the centre line, respectively. **(I, L)** Pairwise comparison of differential gene expression between pre-competent larval (left) and 1 hps postlarval (right) stages. **(I, L)** Pairwise comparison of differential gene expression between 1 hps postlarval (left) and juvenile (right) stages.
