## Supplementary material for "Symbiont biosynthetic output tracks host developmental stage in a marine sponge holobiont": List of Supplementary Tables

Supplementary Table 1. RNASeq Summary: accession details and statistics of raw data for the dual RNASeq holotranscriptome.

Supplementary Table 2. Read counts: normalised mean counts and biological replicate counts for each of the four holobiont partners, with quartiles, DE stats, and complete gene annotations (including Kegg and EC).

Supplementary Table 3. Results of differential gene expression analyses for each of the four holobiont partners, with DE stats and gene annotations.

Supplementary Table 4. Data for heatmaps in Figures 3 and 4 and Supplementary Figure 2: quartiles and mean normalised counts, with gene names and ECs provided.

Supplementary Table 5. *AqS1* enzyme data to support Figure 3F,G: identification of peak stage of expression of *AqS1* genes in Q3 and Q4, with and without an EC designation.

Supplementary Table 6. KEGG BRITE Transporters identified in each of the four holobiont partners, with KO details, DE stats, peak stage of expression, and mean counts provided.

Supplementary Table 7. KEGG PATHWAY Transporters identified in each of the four holobiont partners, with KO details, DE stats, peak stage of expression, and mean counts provided.
